# Human immune system variation during one year

**DOI:** 10.1101/2020.01.22.915025

**Authors:** Tadepally Lakshmikanth, Sayyed Auwn Muhammad, Axel Olin, Yang Chen, Jaromir Mikes, Linn Fagerberg, Anders Gummesson, Göran Bergström, Mathias Uhlen, Petter Brodin

## Abstract

The human immune system varies extensively between individuals, but variation within individuals over time has not been well characterized. Systems-level analyses allow for simultaneous quantification of many interacting immune system components, and the inference of global regulatory principles. Here we present a longitudinal, systems-level analysis in 99 healthy adults, 50 to 65 years of age and sampled every 3^rd^ month during one year. We describe the structure of inter-individual variation and characterize extreme phenotypes along a principal curve. From coordinated measurement fluctuations, we infer relationships between 115 immune cell populations and 750 plasma proteins constituting the blood immune system. While most individuals have stable immune systems, the degree of longitudinal variability is an individual feature. The most variable individuals, in the absence of overt infections, exhibited markers of poor metabolic health suggestive of a functional link between metabolic and immunologic homeostatic regulation.

**HIGHLIGHTS:** Longitudinal variation in immune cell composition during one year

Inter-individual variation can be described along a principal curve

Immune cell and protein relationships are inferred

Variability over time is an individual feature correlating with markers of poor metabolic health

## INTRODUCTION

A well functioning immune system is vital to our health, due to its importance in protecting us from microbial infections, and its dysregulation in many autoimmune and allergic diseases is important. The immune system’s ability to fight cancer has inspired a strong focus on immunotherapy of cancer (Davis and Brodin, 2018). Understanding the composition and function of the human immune system has been challenging due to the high number of inter-dependent cell populations, their complex regulation and the enormous range of variation observed between individuals (Brodin and Davis, 2016). We have learned that immune systems vary as a consequence of both heritable and non heritable influences, the latter exerts a stronger influence on the innate branch of immunity than in the adaptive immune system (Casanova and Abel, 2015; Patin et al., 2018). HLA-genes are exceptions to this rule and provide strong heritable associations with many immune mediated diseases, probably as a consequence of its interactions both with antigen-specific T-cells and innate NK-cells (Horowitz et al., 2013; Matzaraki et al., 2017). Overall, across all cell poulations and plasma proteins, the fraction of variance explained by heritable factors is in the range of 20-40% with a few exceptional cell populations and plama proteins almost entirily explained by heritable traits (Brodin et al., 2015). With the advent of systems-level analyses of many simultaneous cell populations, proteins and mRNA molecules measured in the same sample, the composition and functional potential of human immune systems are becoming more predictable. Examples involve responses to vaccination (Furman et al., 2013; Nakaya et al., 2011; Querec et al., 2008; Sobolev et al., 2016; Tsang et al., 2014), the efficacy of cancer immunotherapy (Krieg et al., 2018), and autoimmune disease processes, which can now be described in more comprehensive ways (Damond et al., 2019; Rao et al., 2017).

The composition of cells in the blood is an individual feature that has been reported to be stable within healthy adults over the course of weeks to months (Carr et al., 2016; Shen-Orr et al., 2016; Tsang et al., 2014). In contrast, very early in life, cell composition changes drastically in response to environmental exposures, suggesting a critical period of development when an individual’s unique cell composition is formed (Olin et al., 2018). In contrast to these reports of stability of adult immune systems, specific cell populations have been reported to change with yearly seasons (Aguirre-Gamboa et al., 2016), and other cell frequencies inferred from blood gene expression patterns, have also been reported to change with the season (Dopico et al., 2015). The overall picture remains unclear. At the same time, the variation in blood cell composition is an important issue to understand given that this is a baseline feature that is predictive of a range of functional responses (Kaczorowski et al., 2017) in vitro, but also in vivo (Tsang et al., 2014). Thus, immune cell composition is a potential useful metric of immunological health. To contribute more data and help settle these uncertainties in human immunology, we report on a longitudinal systems-level analysis performed in 99 healthy individuals, aged 50-65 years, and profiled longitudinally during the course of one year. These individuals are part of the Swedish SciLifeLab SCAPIS Wellness Program (S3WP), and selected from a larger population study, SCAPIS aimed at identifying biomarkers of cardiovascular disease risk among 30,000 Swedes. In these individuals, we find a broad range of variation in immune cell composition which can be effectively visualized along a principal curve with extreme phenotypes readily identified. We find that fluctuations over time occur, that cannot be explained by season, but can be used to infer cell and protein relationships from co-regulated features. At the global level, over-time immune variation is an individual feature with the most variable individuals also exhibiting markers of poor metabolic health suggestive of a loss of homeostatic regulation.

## RESULTS

### Longitudinal profiling of healthy adults during one year

Within the national cohort study SCAPIS, 30,000 clinically healthy and middle age Swedes are enrolled and subject to deep cardiovascular phenotyping. From this cohort, a 101 individual subset was enrolled to undergo deep molecular phenotyping within the S3WP program at the Science for Life Laboratory (Figure 1A). 99 of these 101 individuals returned for all four visits during the course of one year (Figure 1B), viable peripheral blood mononuclear cells (PBMCs) and plasma samples were frozen and stored until analysis. We analyzed around 200,000 cells per sample, and 78,891,056 cells in total by mass cytometry using a 38-parameter panel targeting all mononuclear immune cell populations. In plasma samples, 11-panels of Olink assays were applied, capturing 978 proteins in total. Out of these, 750 plasma proteins with highest confidence of detection was retained for downstream analyses (Experimental Procedures). The 101 subjects enrolled, were 53 women and 48 men, aged 50-65 years (Figure 1C), with variable in body-mass index and blood pressure levels (Figure 1D-E).

**Figure 1.**
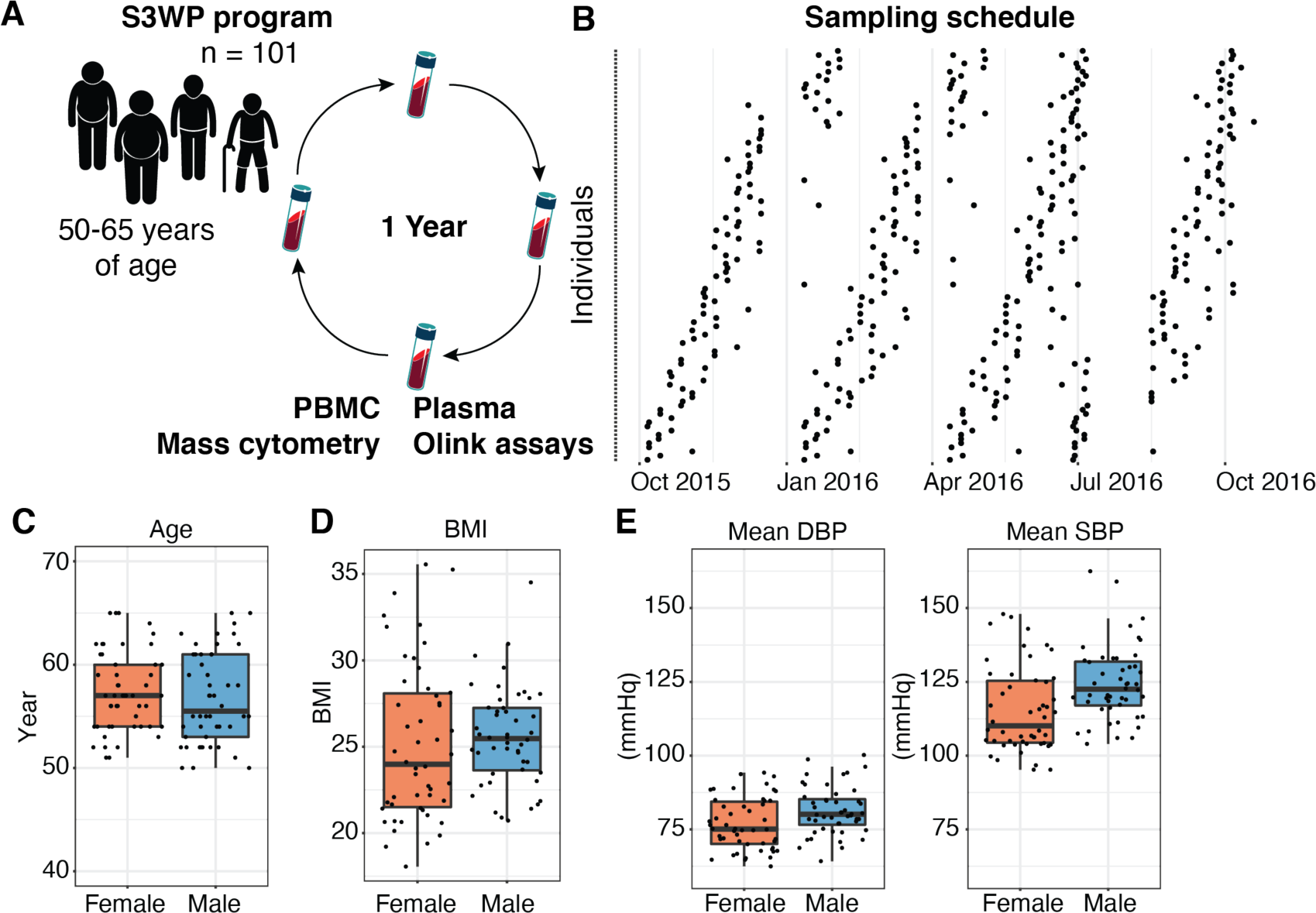
101 healthy adults, profiled longitudinally during one year. **(A)** Samples collected approximately every third month during one year were analyzed by Olink assays (978 Plasma proteins), and Mass cytometry (PBMC cell populations). **(B)** individuals (rows) and their respective sampling times during 2015 and 2016 are shown. **(C)** Age, **(D)** Body Mass Index, BMI, and **(E)** blood pressure distributions for female (n=53) and male (n=48) subjects in the cohort respectively.

### Describing immune variation along a principal curve

A comprehensive analysis of immune system variation requires the simultaneous analysis of many cell populations. Because each of these vary substantially among individuals, their collective frequencies are better metrics for accurately describing immune variation than each of the cell frequencies individually (Kaczorowski et al., 2017). We manually gated 115 immune cell populations in each sample, but because these are hierarchical in nature, we only included 53 non-redundant populations for compositional analyses. In this dataset all cell population frequencies have a sum equal to 1. To describe variation among 396 samples from 99 individuals with four samples each, we calculated an Aitchison’s distance matrix (Templ et al., 2011), and visualize samples by multidimensional scaling (MDS) (Figure 2A). Here, each sample is positioned based on its pairwise distances to all other samples. The distribution of individual samples is continuous, without any evidence for discrete “immunotypes” within the cohort (Kaczorowski et al., 2017) (Figure 2A).

**Figure 2.**
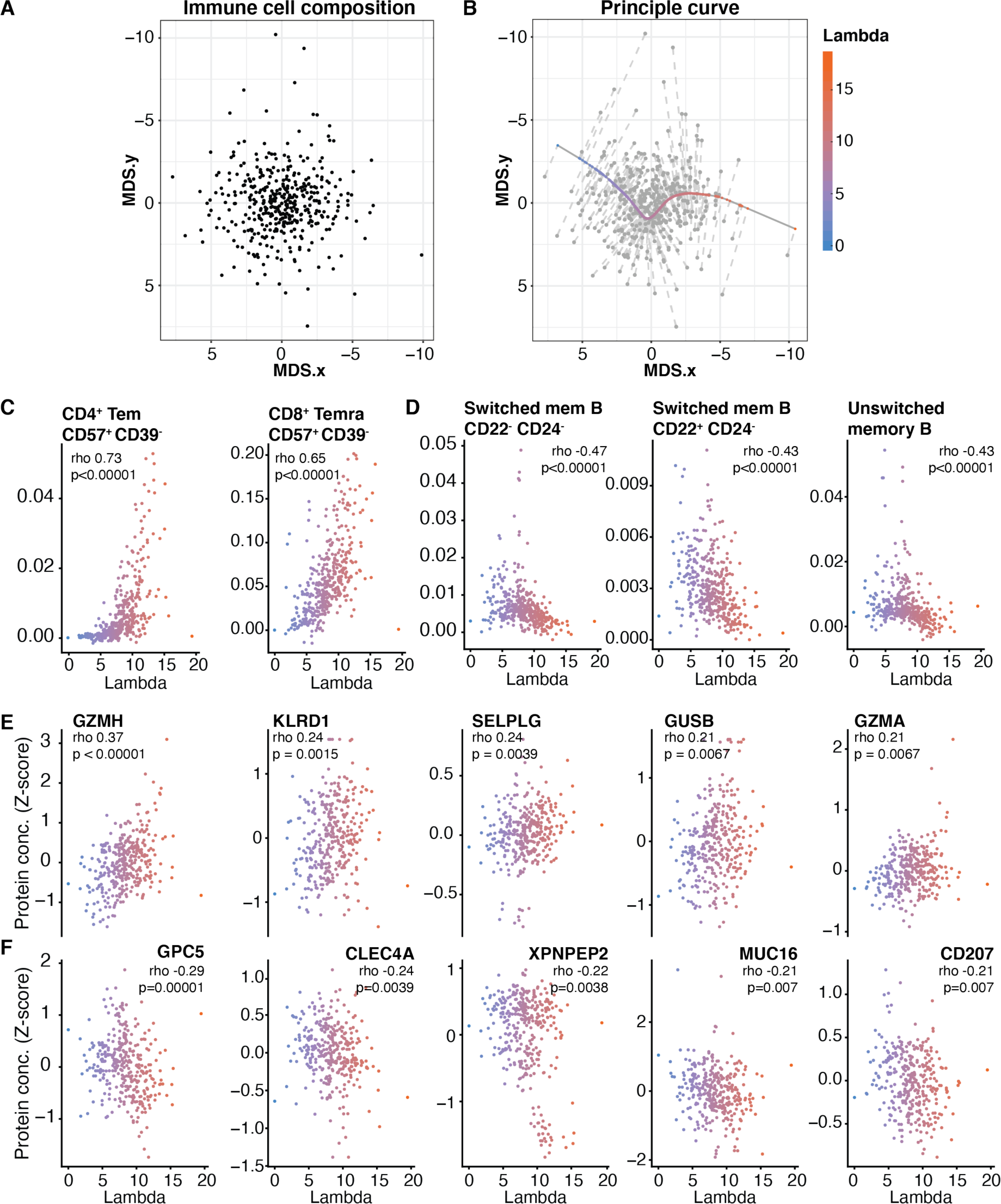
Describing variation in global immune cell composition along a principal curve. **(A)** Pairwise distances (Aitchison’s) based on relative abundances of 53 non-overlapping immune cell populations. **(B)** A principal curve is shown that captures the variance in immune cell composition (of 53 non-redundant cell populations) in 396 samples obtained from 99 individuals. Each dot represents each individual sample. **(C)** Two of the most positively correlated cell populations increasing along the principal curve. **(D)** Three of the most negatively correlated cell populations decreasing along the principal curve. **(E)** Five of the most positively correlated plasma proteins, increasing along the principal curve. **(F)** Five of the most negatively correlated plasma proteins decreasing along the principal curve.

To summarize the range of variation using a simpler metric, we fit a principal curve that best explain the variation in immune cell composition (Experimental Procedures)(Figure 2B). This Principal curve is the smooth curve that pass through the middle of the data cloud and is used as a simplified representation of the variation in the data (Hastie and Stuetzle, 1989). Each individual sample is later projected onto this principal curve, and their location along this line is denoted by a parameter lambda (Figure 2B). In this way, lambda serves as a simplified metric of variation in immune cell composition. The main benefit of using such a measure of variation is that investigations of individual cell populations along this range of variation, and investigations of extreme phenotypes at the ends of the lambda spectrum are made possible (Figure 2B). We calculated correlations (Spearman) between individual cell frequencies and the lambda metric and found 38 immune cell populations correlated with lambda (p<0.05). The strongest positive correlations was seen for CD4^+^ effector memory and CD8^+^ T-EMRA populations (Figure 2C), and class-switched and un-switched memory B-cell populations that were negatively correlated with lambda (Figure 2D). We conclude that a principal curve can describe global immune variation and reveal relationships such as the opposing relationship between effector T- and memory B-cell populations in healthy adults. By projecting additional samples onto the principal curve, immune system differences among individuals with different clinical outcomes or therapeutic responses will be possible in the future.

In the analysis above, only immune cell frequencies are considered when defining the lambda metric, but this value can also be used to assess relationships between global immune cell composition and plasma protein concentrations. To this end, we correlated abundances of 750 plasma proteins measured by Olink assays (Olink AB, Uppsala)(Experimental Procedures) to lambda. This resulted in 23 significant correlations (p<0.05) with Granzyme H, and A, KLRD1, SELPLG and GUSB being the strongest positive ones (Figure 2E), and GPC5, CLEC4A, XPNPEP2, MUC16 and CD207 as the strongest negatively correlated proteins along lambda (Figure 2F). The correlation (Spearman) coefficients with lambda for all cell populations and plasma proteins is also available online (Supplementary Table 1). These results suggest that global immune cell composition can be predicted from a smaller number of plasma protein biomarkers, although this will need to be validated further. We conclude that immune variation in humans is not random, but determined by specific cell and protein relationships that can be described by a simplified metric allowing systems-level comparisons among patients, conditions and in relation to therapeutic interventions to be made.

### Sex-differences in global immune cell composition

Autoimmune diseases segregate largely by sex, but the reasons for these differences are poorly understood (Ngo et al., 2014). We compared global immune cell compositions among the 49 men and 52 women in our cohort plotted pairwise Aitchison’s distances among samples by MDS (Figure 3A). We calculate differences between the sexes using a permutation analysis of variance (Permanova) (Templ et al., 2011), which show a significant difference between male and female sample distributions (p<0.001), but a limited amount of variance explained (sums of squares). Sex explained only 2.39% of the total variance in cell composition (Figure 3A). The most differing cell populations between men and women were naïve CD4^+^ T-cells, higher in females while activated and antigen experienced CD8^+^ T-cells were more abundant in male subjects (Figure 3B). We conclude that global cell composition, like some individual cell populations reported previously (Aguirre-Gamboa et al., 2016; Carr et al., 2016; Patin et al., 2018), differ between men and women, but explaining only a minor fraction of the overall variance (Aguirre-Gamboa et al., 2016; Carr et al., 2016; Patin et al., 2018).

**Figure 3.**
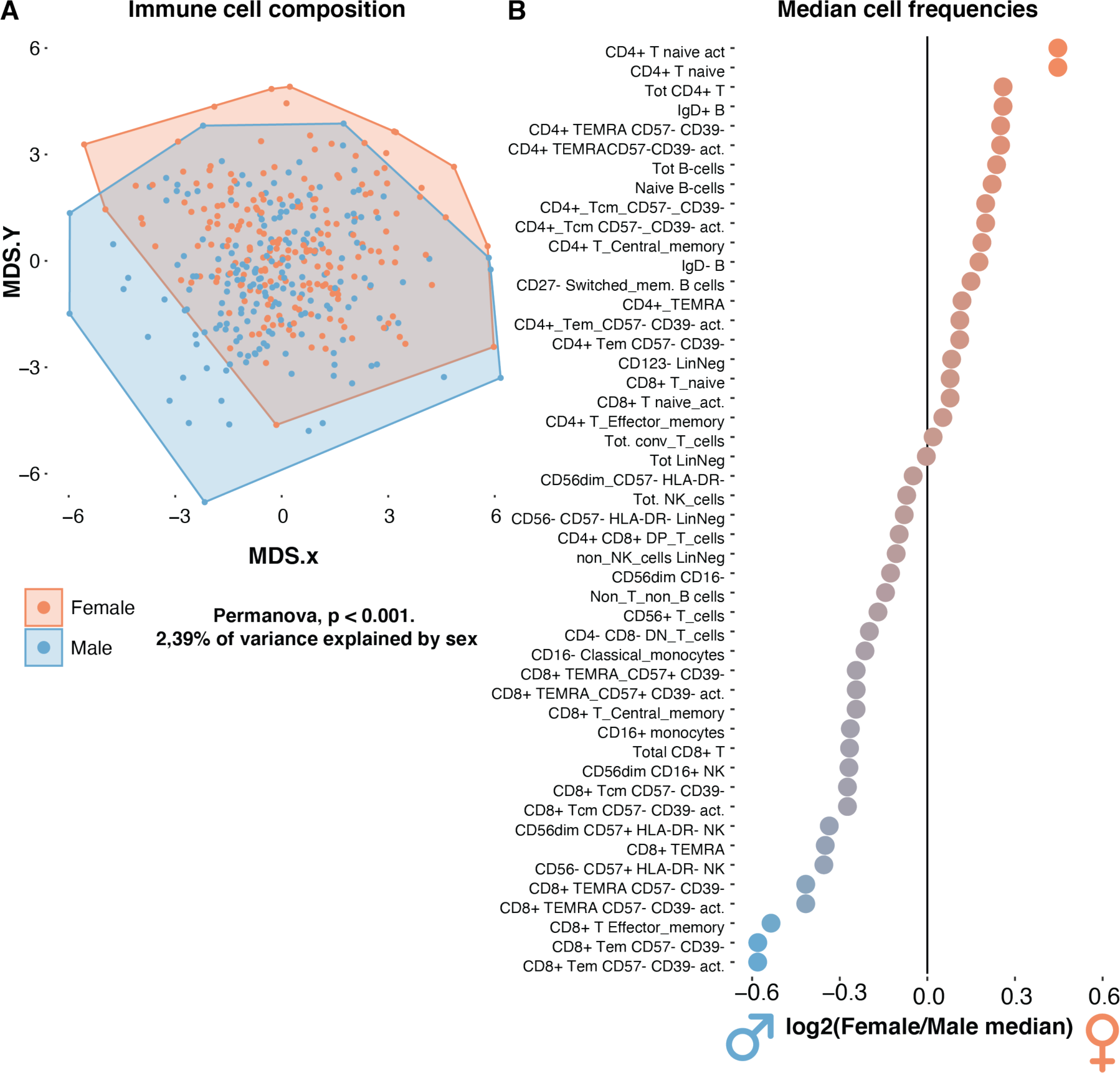
Sex-differences in cell composition. **(A)** Pairwise distances (Aitchison’s) based on relative abundances of 53 non-overlapping immune cell populations visualized by MDS in 396 samples from 53 women and 48 men. **(B)** 48 non-zero immune cell population frequencies as log_2_ ratios (Female/Male).

### Global immune cell composition is stable over the seasons

Another important factor previously shown to influence individual immune cell populations (Aguirre-Gamboa et al., 2016) and gene expression (Dopico et al., 2015) is seasonality. Based on the lambda metric of global immune cell composition shown in Figure 2B for each sample, the overall observation is that global immune cell composition is stable over the course of the 12 months of a year (Figure 4A). In fact there were no significant shifts in global immune cell composition between any of these months, suggesting that overall immune variance is not strongly influenced by season (Figure 4A).

**Figure 4.**
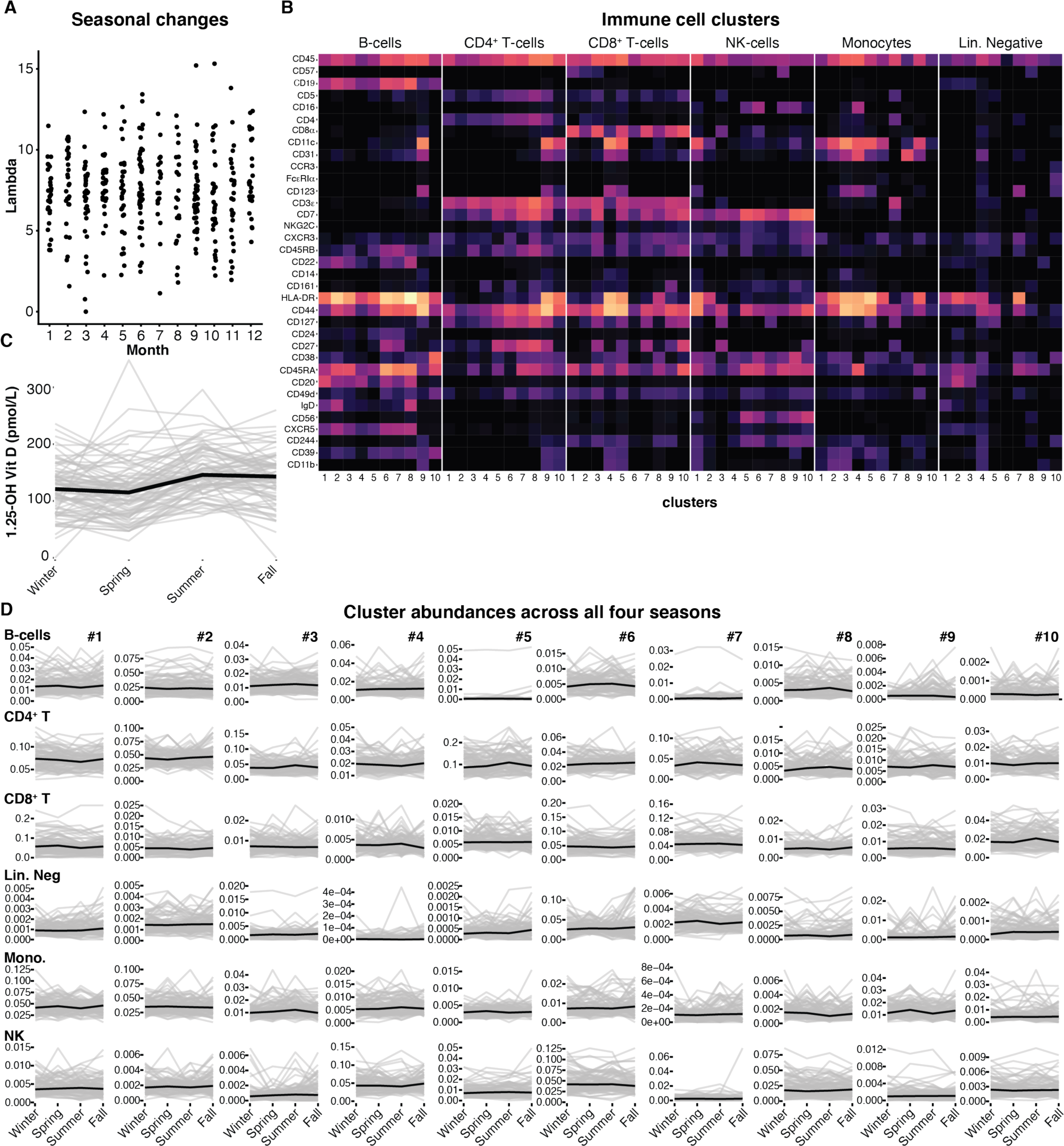
Stable immune cell frequencies across seasons. (**A**) Lambda values as shown in Figure 2, for samples collected at the indicated month. (**B**) Immune cell clusters defined within the indicated cell lineages using Self-organizing map, SOM clustering. Clusters in columns and their respective marker expression in rows. (**C**) Serum levels of 1.25-OH Vitamin D across individuals (n=82), with one sample taken per season; Winter (Dec-Feb), Spring (March - May), Summer (June - August), and Autumn (September - November). Grey lines show individual level data and black line represent mean concentration. (**D**) Immune cell cluster abundances within individuals sampled every season (n=81), Winter (Dec-Feb), Spring (March - May), Summer (June - August), and Autumn (September - November). Grey lines show individual level data and black line represent mean abundance of the indicated cell cluster.

One important limitation of previous analyses of seasonal variation in immunity is that these involved comparisons across different individuals, each sampled at different months of the year, rather than analyses within the same individuals sampled longitudinally. Here we have taken advantage of the longitudinal samples collected every three months during the course of one year. To investigate seasonal changes involving specific subpopulations of cells, self-organizing maps were used to cluster subpopulations of cells, within the major CD4^+^ T-, CD8^+^ T-, B-, and NK-cell lineages, as well as Monocytes, and a remaining lineage negative population (Figure 4B). This clustering approach allows for previously undefined subpopulations with possible seasonal fluctuations to be identified, while manual gating of known populations might miss such seasonal populations. To allow comparisons among individuals, even when the individuals are sampled at slightly different months of the year, we grouped samples according to the four weather seasons in Sweden; Winter (Dec-Feb), Spring (March-May), Summer (June-August), and Autumn (September-November). To verify that these seasons capture known changes during the year in Sweden, we show 1.25-dihydroxy Vitamin D-levels for 83 individuals with such a measurement taken once in every season (Figure 4C). We find higher concentrations of vitamin D in most individuals during the summer and autumn due to the longer days and increased exposure to sunlight during these seasons in Sweden (Klingberg et al., 2015). In the cohort, 81 individuals also had one immune cell measurement by mass cytometry performed during each of these four seasons, allowing us to investigate seasonal fluctuations in cluster abundances (Figure 4D). We find large inter-individual variation in clusters abundances, and for some clusters also some strong variation over time in selected individuals, but this longitudinal variation did not follow seasonal patterns shared by many individuals (Figure 4D).

### Cell and protein relationships inferred from coordinated fluctuations over time

Although no reproducible seasonal patterns were seen here, we decided to investigate the longitudinal variability of immune cell abundances further. In particular, we were interested to take advantage of fluctuations in cells and plasma proteins, to identify co-regulated immune system components, and in particular dependencies between plasma proteins and cell frequencies (Figure 5A). The 750 most reliably detected plasma proteins, and 115 manually gated immune cell populations represent an enormous number of possible associations. To be able to interpret such associations, a more manageable number of features is required. We therefore focused on 32 canonical immune cell populations, and the 378 proteins that best represents the overall variance in the plasma protein dataset (Experimental procedures). Using a Bayesian variable selection method, sets of proteins are selected randomly as predictors of a given cell population’s abundance in a regression model (Carbonetto and Stephens, 2012). When performed iteratively, the posterior probability of inclusion (ppi) represents a measure for the likelihood of association between a protein and cell population (Experimental procedures). By thresholding associations from this iterative variable selection procedure (Supplementary Figure 1), resulted in a network model of the 226 significant associations (FDR = 0.01), out of 12,096 possible associations (Supplementary Table 2). Out of these 106 were positive associations (positive beta coefficient), and 120 negative associations (negative beta coefficients). The 50 strongest positive, and 50 strongest negative associations are shown in Figure 5B.

**Figure 5.**
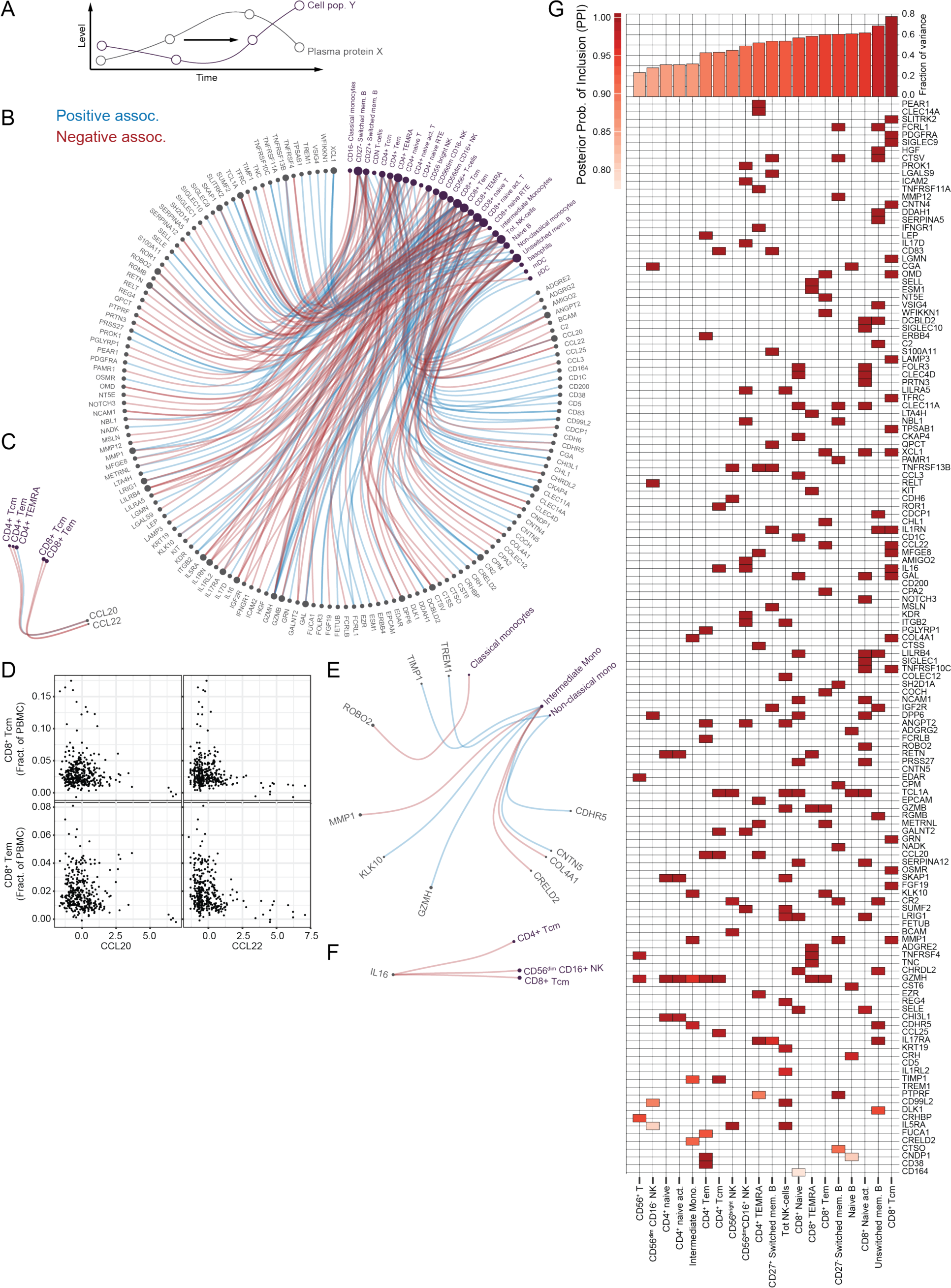
Inference of cell - protein relationships. **(A)** Coordinated changes in individual cell populations and plasma protein levels allow for inference of inter-dependencies using a Bayesian variable selection method. **(B)** Top 50 positive (blue) and the 50 strongest negative (red) associations, between 32 core cell populations and 378 proteins explaining most of the variance. Node size is proportional to the number of associations. **(C)** Highlight of associations between chemokines CCL20 and CCL22 and 5 related T-cell subsets. (D) Dot plot showing individual sample measurements of CCL20, CCL22 vs. CD8^+^ Tcm and CD8^+^ Tem respectively. **(E)** Negative association between the cytokine IL-16 and central memory T-cells and CD56dim NK cells. **(F)** Different protein associations for different monocyte subpopulations. **(G)** variance in abundance of each indicated cell population (columns), collectively explained by the indicated plasma proteins (rows).

Some examples involve the chemokines CCL20 signaling via the CCR6 receptor (Schutyser et al., 2000), and CCL22 binding CCR4 (Vulcano et al., 2001), both negatively associated with central memory and effector memory population frequencies, in both CD4^+^ and CD8^+^ T-cell lineages (Figure 5C). These negative associations are driven by extreme phenotypes because subjects with very high levels of these cell populations are always low in plasma CCL20 and CCL22, and vice versa (Figure 5D).

Another example involves the lesser known cytokine IL-16 which signals directly via the CD4 receptor on T-cells, and potentially on other cells expressing this receptor. IL-16 has also has been shown to influence the interaction of CD4^+^ T-cells with B-cells, DCs and importantly other T-cell populations, suggesting a more broad modulatory role (Skundric et al., 2015). We found strong negative associations between plasma IL-16 levels and central memory T-cells, both CD4^+^ and CD8^+^, as well as cytotoxic CD56^dim^ NK cells, cells that are known to express CD4 when activated (Bernstein et al., 2006)(Figure 5E).

In contrast to the shared protein associations for central memory T-cells, the three main monocyte populations are associated with different plasma proteins, suggesting that modulation of specific subpopulations might be possible (Figure 5F). These examples highlight novel hypotheses that can be generated from longitudinal analyses of co-regulated immune cells and proteins. Finally, we wanted to assess how much of the total variance in cell population abundance, that could be attributed to specific plasma proteins. We found that nearly 80% of the variance in central memory T-cell abundance across samples could be explained by the concentration of 21 plasma proteins, while a more marginal set of only 4 proteins associated with CD56^+^ T-cells, explained >20% of its variance (Figure 5G). Specific plasma proteins were more informative than others, such as Granzyme H, (GZMH) found to be associated with 8 of the 20 most predictable populations (>20% of variance explained)(Figure 5G). This analysis would enable smaller sets of highly informative measurements to be validated in larger cohorts and possibly making human immune systems more predictable in the future.

### Immune variation over time is an individual feature

Next we wanted to investigate the degree of variation over time, not at the level of individual immune system components, but at the level of the global cell composition. Such profiles have previously been reported to be stable in healthy adults over the course of weeks to months (Carr et al., 2016; Shen-Orr et al., 2016; Tsang et al., 2014), while young children change drastically, before an individual’s stable phenotype has been established (Olin et al., 2018). In our cohort, we found that samples from the same individual were more similar (Aitchison’s distance), as compared to samples from different individuals (Figure 6A). Using the same distance metric, comparing global cell composition between two samples, we calculated the total and mean distances travelled across all four study visits (Figure 6B-C). This metric can be used to compare individuals with respect to their overall variability during the year of study. A few individuals reported infections between visits and this was typically associated with individual visits differing largely from the mean in a few subjects but did not affect the overall variability across all four visits (Figure 6C). The general pattern however was that most individuals had stable immune cell composition over time, but with a degree of variability that is an individual feature, not clearly explained by reported infectious disease events, but rather a feature of their immune systems giving rise to larger shifts in phenotype between every timepoint during the year (Figure 6C).

**Figure 6.**
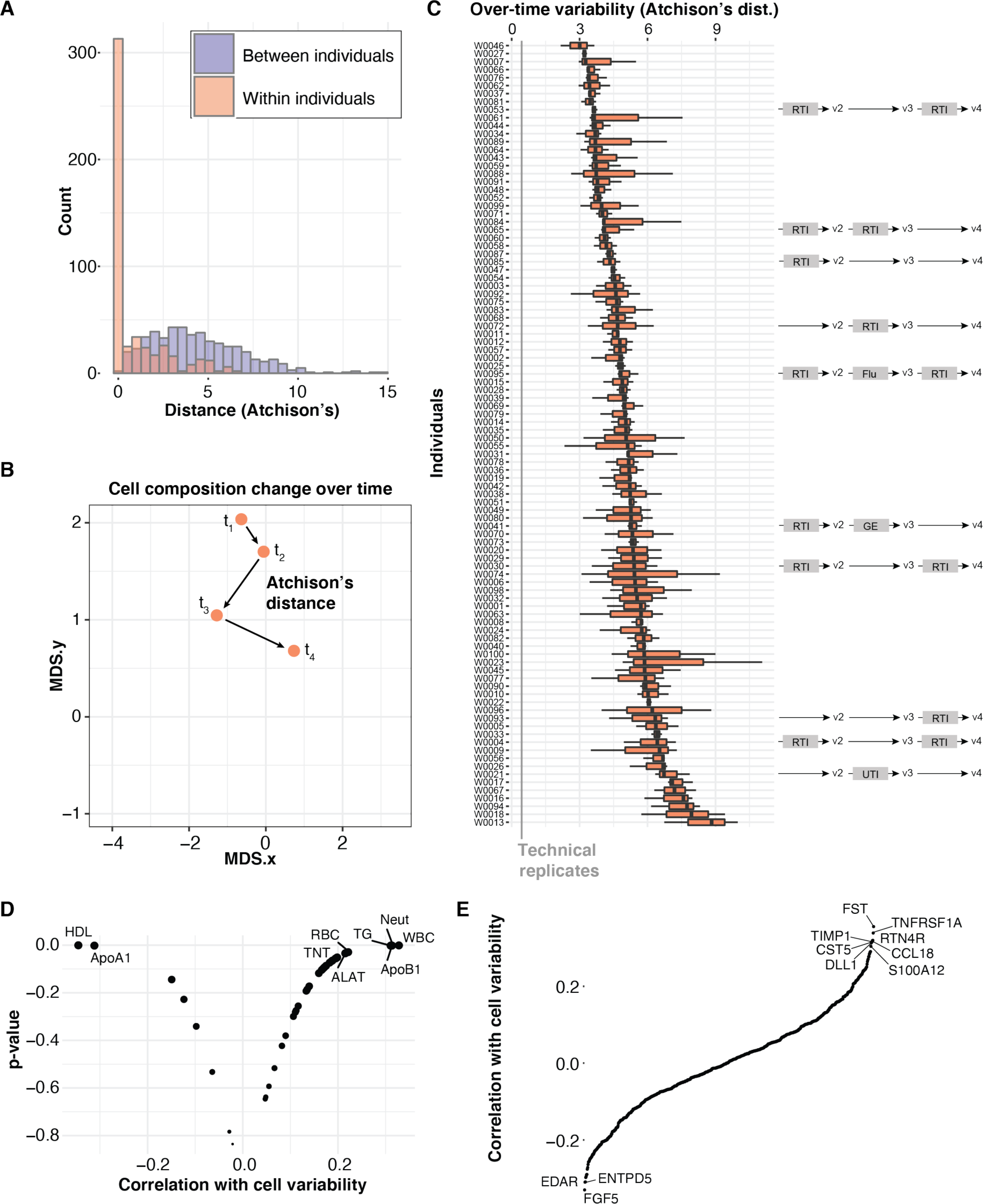
Over-time variability is an individual feature associated with poor metabolic health. **(A)** Aitchison’s distances calculated between samples taking all collective cell frequencies into account. Distances between multiple samples within an individual (Orange) and distances between samples from different individuals (Purple) shown as separate bar charts. **(B)** Illustration of distances travelled by one individual across 4 time-points in MDS space. **(C)** Intra-individual distances (Aitchison’s) shown for all individuals ordered from low to high mean distance travelled. Grey line indicate Aitchison’s distances across technical replicates and time-lines on the right indicate self-reported infections; UTI: Urinary Tract Infection, RTI: Respiratory Tract Infection, GE: Gastroenteritis. **(D)** Volcano plot of clinical measurements (mean values) positively and negatively correlated (Spearman) with over-time variability in immune cell composition. Significantly correlated features (p<0.05) are labeled. (E) Plasma proteins ranked by their correlation (Spearman), with over-time variability. Proteins with rho > 0.3 or < 0.3 are indicated.

### High-variability is associated with markers of poor metabolic health

We next wanted to understand what mediates this individual degree of variation, expressed as a different mean distance travelled between samplings. There was no link between time in-between visits and degree of variation (Data not shown). There were also no significant correlations (Spearman, FDR< 1%), between plasma proteins and variability. We then investigated relationships between clinical chemistry measurements at each visit and the degree of variability. As shown in Figure 6D, we found a positive correlation between multiple markers of poor metabolic health, such as plasma Apo-B, Triglycerides (TG), and LDL cholesterol, and conversely a negative association between over-time immune variability (mean Aitchison’s distance), and markers of metabolic health, such as HDL, and Apo-A1 (Figure 6D). Also, highly-sensitive Troponin T, a heart muscle leakage protein and a marker of cardiovascular disease, was positively correlated with immune variability (Figure 6D). Moreover, the total white blood cell count, red blood cell count, RBC and neutrophil count were also positively associated with high immune variability over time (Figure 6D). When looking at plasma proteins correlated with over-time cell variability, CCL18 was one of the top hits and this chemokine has previously been shown to be strongly expressed by macrophages within atherosclerotic lesions (Hägg et al., 2009) and correlated with the extent of coronary artery disease in human patients (Versteylen et al., 2016).

These findings suggest that immune system variability is an individual feature, not explained by overt responses to infections, but instead associated with markers of poor metabolic health. Although, such associations must be validated in additional cohorts and follow-up in mechanistic analyses these data suggest that over time variability is an important feature to consider in human immunology. Longitudinal, systems-level data thereby allowed for a more precise description of inter-individual variation, the inference of biological associations between cells and proteins and a proposal for variability as a novel feature, indicative of immune system health and regulation, and associated with metabolic and cardiovascular health and homeostasis.

## DISCUSSION

In this manuscript we have performed a longitudinal analysis of clinically healthy middle-aged individuals over the course of one year. We find that variation can be described along a principal curve and a lambda parameter defined which can then more easily be analyzed in relation to disease manifestations, metadata or other parameters of interest. We also show the potential of using longitudinal data in humans as a means of learning inter-dependencies among hundreds of immune system components. We also assessed the variability over time in relation to season without finding any cell populations that were strongly associated with season. This finding contrasts some previous analyses which have suggested such seasonal changes (Aguirre-Gamboa et al., 2016; Dopico et al., 2015). Our study differs from these in that we assessed seasonal changes from longitudinal samples within a given individual, while these previous analyses involved different individuals sampled once at different time-points during the year. We believe our study design is more appropriate for directly assessing seasonal changes, although having multiple samples across multiple consecutive years would have been even stronger for detecting seasonal changes reoccurring over consecutive years. Given our previous work on immune variation in monozygotic twins (Brodin et al., 2015), one possibility would be to monitor and compare such MZ twins longitudinally across seasons to further dissect possible seasonal changes occurring in a reproducible fashion in co-twins.

In a previous report we showed that newborn immune systems change drastically in their cell composition, as a response to environmental exposures after birth (Olin et al., 2018). In contrast, adult immune systems have previously been shown to be stable over the course of weeks, months and even a few years (Shen-Orr et al., 2016). At the same time, immune system composition in older individuals is more heterogeneous as compared to younger individuals (Kaczorowski et al., 2017) and a recent analysis even use such variability to estimate immunological age that is predictive of all-cause mortality and cardiovascular health (Alpert et al., 2019). Monozygotic twins diverge over time, suggesting a cumulative influence of environmental factors over the course of life shaping human immune systems. Here we show that variability over time in middle-aged individuals is actually an individual feature, even in the absence of overt infections or inflammatory conditions. The finding that high variability is associated with markers of cardiovascular disease risk and poor metabolic health, suggests that over time variability might be also be an interesting biomarker of poor immunological health that should be studied in relation to disease. One particular condition that comes to mind here is “inflammaging”, a chronic inflammatory state in some individuals in older age, with increased mortality, and sharing many features with the inflammation seen in obesity and nutrient excess (Franceschi et al., 2018). Perhaps, increased immune variability and loss of immune homeostasis is a feature of both such chronic inflammatory states.

One limitation of the current study is that the population studied is relatively small and not representative of all human populations. For example the HLA-diversity of human populations cannot be accounted for using a population of this size. Also, in a different population living in a different environmental condition, and facing a different disease panorama a different pattern of variability over time might be seen. In a parallel study involving this same S3WP consortium the longitudinal variability of the gut microbiome has been assessed, but we found no correlation between immune system variability and gut microbiome variability (Data not shown). This is interesting given that the variation in gut microbiome composition has previously been reported to also be an individual feature (Flores et al., 2014), but presumably under the influence of other factors as compared to the immune cell composition described herein. One important factor not accounted for here is Cytomegalovirus, (CMV) previously shown to affect immune systems broadly (Brodin et al., 2015; Kaczorowski et al., 2017; Piasecka et al., 2018), but due to ethical permit limitations not available for study in the S3WP cohort.

## ACKNOWLEDGEMENTS

We would like to thank all S3WP program coordinators, sample collectors, volunteers and data collectors within the consortium. Also, we would like to thank all members of the Brodin lab and several members of the S3WP program for inspiring discussions. The study is supported by a grant from the Erling Persson Foundation to M.U, and grants to P.B from Karolinska Institutet, Science for Life Laboratory, and the Swedish research council. The larger SCAPIS study from which subjects are enrolled, is supported by the Swedish Heart and Lung Foundation, the Knut and Alice Wallenberg Foundation, the Swedish Research Council and VINNOVA.

---

**Suppl. Fig 1.**
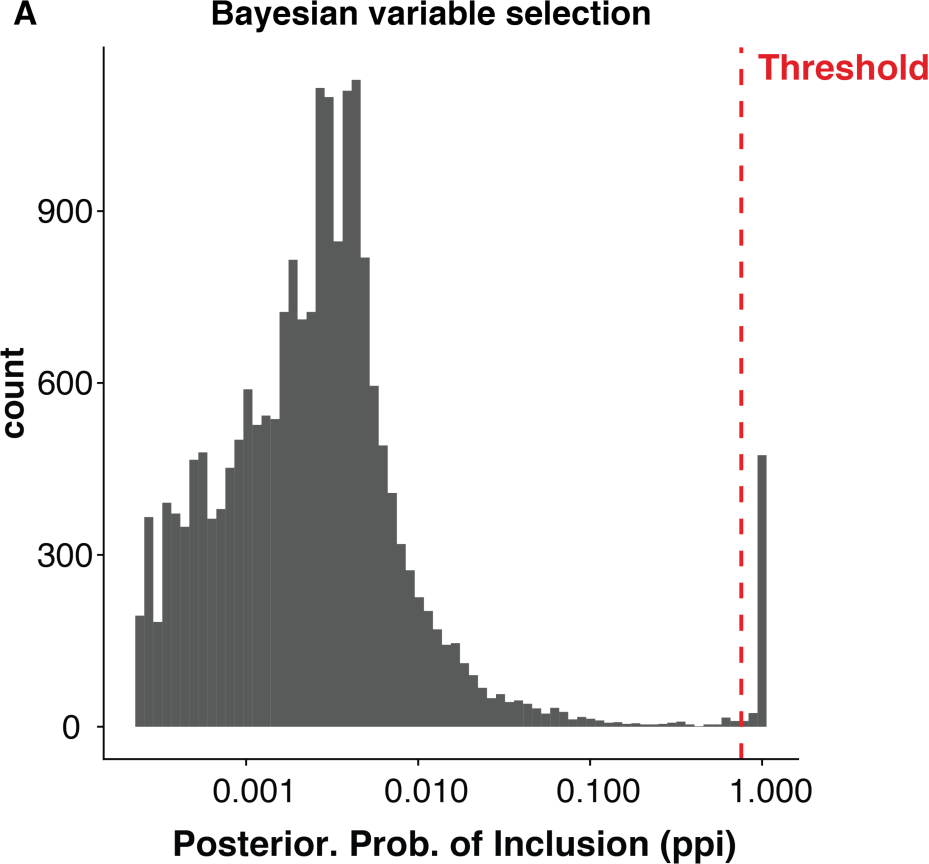
Distribution of posterior probabilities of inclusion and the threshold for inclusion in the analysis (red line). Related to figure. 5.

## STAR METHODS

### KEY RESOURCES TABLE

KEY RESOURCES TABLE

| REAGENT or RESOURCE | SOURCE | IDENTIFIER |
| --- | --- | --- |
| **Antibodies** |  |  |
| CD45 (HI30) – 89Y | Fluidigm | Cat# 3089003B; RRID:2661851 |
| CD57 (HCD57) – Purified | BioLegend | Cat# 322302; RRID:AB_535988 |
| CCR6 (11A9) – Purified | BD | Cat# 559560; RID:AB_2291135 |
| CD19 (HIB19) – 142Nd | Fluidigm | Cat# 3142001B; RRID:AB_2651155 |
| CD5 (UCHT2) – Purified | BioLegend | Cat# 300602; RRID:AB_314088 |
| CD16 (3G8) – Purified | BioLegend | Cat# 302002; RRID:AB_314202 |
| CD4 (RPA-T4) – 145Nd | Fluidigm | Cat# 3145001B; RRID:AB_2661789 |
| CD8a (SK1) – Purified | BioLegend | Cat# 344702; RRID:AB_1877104 |
| CD11c (Bu15) – 147Sm | Fluidigm | Cat# 3147008B; RRID:AB_2687850 |
| CD31 (WM59) – Purified | BioLegend | Cat# 303102; RRID:AB_314328 |
| CD193/CCR3 (5E8) – Purified | BioLegend | Cat# 310702; RRID:AB_345394 |
| FcεRIα (AER-37) – Purified | BioLegend | Cat# 334602; RRID:AB_1227649 |
| CD123 (6H6) – Purified | BioLegend | Cat# 306002; RRID:AB_314576 |
| TCRαβ (IP26) – Purified | BioLegend | Cat# 306702; RRID: AB_314628 |
| CD3e (UCHT1) – 154Sm | Fluidigm | Cat# 3154003B; RRID:AB_2687853 |
| CD7 (CD7-6B7) – Purified | BioLegend | Cat# 343102; RRID:AB_1659214 |
| NKG2C (134591) – Purified | R&D | Cat# MAB138; RRID:AB_2132982 |
| CXCR3 (G025H7) – Purified | BioLegend | Cat# 353733; RRID: AB_2563724 |
| CD45RB (MEM-55) – Purified | BioLegend | Cat# 310202; RRID:AB_314805 |
| CD22 (HIB22) – Purified | BioLegend | Cat# 302502; RRID:AB_2074593 |
| CD14 (M5E2) – Purified | BioLegend | Cat# 301802; RRID:AB_314184 |
| CD161 (HP-3G10) – Purified | BioLegend | Cat# 339902; RRID:AB_2661837 |
| HLA-DR (L243) – Purified | BioLegend | Cat# 307602; RRID:AB_314680 |
| CD44 (BJ18) – Purified | BioLegend | Cat# 338802; RRID:AB_1501199 |
| CD127 (A019D5) – 165Ho | Fluidigm | Cat# 3166007B; RRID:AB_2661803 |
| CD24 (ML5) – Purified | BioLegend | Cat# 311102; RRID:AB_314851 |
| CD27 (L128) – 167Er | Fluidigm | Cat# 3167006B; RRID: AB_2687645 |
| CD38 (HIT2) – Purified | Biolegend | Cat# 303502; RRID:AB_314354 |
| CD45RA (HI100) – 169Tm | Fluidigm | Cat# 3169008B |
| CD20 (2H7) – Purified | BioLegend | Cat# 302302; RRID:AB_314250 |
| CD49d (9F10) – Purified | BioLegend | Cat# 304302; RRID:AB_314428 |
| IgD (IA6-2) – Purified | BioLegend | Cat# 348235; RRID:AB_2563775 |
| CD56 (NCAM16.2) – Purified | BD | Cat# 559043; RRID:AB_397180 |
| CD185/CXCR5 (51505) – Purified | R&D | Cat# MAB190; RRID:2292654 |
| CD244/2B4 (C1.7) – Purified | BioLegend | Cat# 329502; RRID:1279194 |
| CD39 (A1) – Purified | BioLegend | Cat# 328202; RRID: 940438 |
| CD11B (Mac-1) – 209Bi | Fluidigm | Cat# 3209003B; RRID:AB_2687654 |
| **Biological Samples** |  |  |
| Healthy subjects from S3WP | University of Gothenburg | N/A |
| **Chemicals, Peptides, and Recombinant Proteins** |  |  |
| RPMI 1640 medium | Sigma-Aldrich | Cat# R848 |
| FBS | Sigma-Aldrich | Cat# 12103C |
| Penicillin-streptomycin | Sigma-Aldrich | Cat# P4333 |
| Benzonase | Sigma-Aldrich | Cat# E1014 |
| 1X PBS | Rockland | Cat# MB-008 |
| EDTA | Rockland | Cat# MB-014 |
| Sodium Azide | Sigma-Aldrich | Cat# 71289 |
| Bovine Serum Albumin | Sigma-Aldrich | Cat# A3059 |
| Paraformaldehyde | Polysciences | Cat# 00380-1 |
| Intercalator-Ir | Fluidigm | Cat# 201192B |
| EQ Four Element Calibration Beads | Fluidigm | Cat# 201078 |
| Maxpar X8 Multimetal labeling kit | Fluidigm | Cat# 201300 |
| Metal isotopes as chloride salts | Fluidigm | N/A |
| Dimethyl Sulfoxide | Sigma-Aldrich | Cat# D8418 |
| Cell-ID Cisplatin | Fluidigm | Cat# 201064 |
| Metal isotopes as chloride salts | Trace Sciences International | N/A |
| Protein Stabilizer PBS | Candor Bioscience GmbH | Cat# 131125 |
| Critical Commercial Assays |  |  |
| Test tube with 35 μm nylon mesh | Corning | Cat# 352235 |
| Cardiometabolic panel | Olink AB | N/A |
| Cell Regulation panel | Olink AB | N/A |
| Cardiovascular II panel (CVD II) | Olink AB | N/A |
| Cardiovascular III panel (CVD III) | Olink AB | N/A |
| Development panel | Olink AB | N/A |
| Immune Response panel | Olink AB | N/A |
| Immuno-Oncology panel | Olink AB | N/A |
| Oncology II panel | Olink AB | N/A |
| Inflammation panel | Olink AB | N/A |
| Metabolism panel | Olink AB | N/A |
| Neurology, and Organ Damage panel | Olink AB | N/A |
| Deposited Data |  |  |
| FCS files, Mass cytometry | This paper |  |
| Protein expression data | This paper |  |
| Software and Algorithms |  |  |
| Mass Cytometry Normalizer | Finck et al., 2013 | https://github.com/nolanlab/bead-normalization/releases |
| CyTOF software (v. 6.0.626) | - | https://www.fluidigm.com/ |
| robCompositions 2.0.5 | Templs et al., 2011 | https://cran.r-project.org/web/packages/robCompositions/index.html |
| Kohonen 3.0.2 | Wehrens and Buydens, 2007 | https://cran.r-project.org/web/packages/kohonen/index.html |
| dmbvs | Wadsworth et al., 2017 | https://github.com/duncanwadsworth/dmbvs |
| Princurve 2.1.3. | Hastie and Stuetzle, 1989 | https://cran.r-project.org/web/packages/princurve/index.html |
| MASS 7.3-51.1 | Venables and Ripley, 2002 | https://cran.r-project.org/web/packages/MASS/index.html |
| Sva 3.18.0 | N/A | http://bioconductor.org/packages/release/bioc/html/sva.html |

### CONTACT FOR REAGENT AND RESOURCE SHARING

Further information and requests for resources and reagents should be directed to Petter Brodin. Raw FCS files and plasma protein levels will be shared to anyone after certification that data will not be further disseminated to any third party. This mechanism is not the choice of the authors but the restriction imposed by the legal department, Gothenburg University who is responsible for data management within the SCAPIS project.

### EXPERIMENTAL MODEL AND SUBJECT DETAILS

#### Wellness subjects

From an ongoing Swedish SciLifeLab’s SCAPIS (**S**wedish **CA**rdio**P**ulmonary bio**I**mage **S**tudy) Wellness Profiling (S3WP) program, which is a prospective observational study of a randomly selected 30,000 Swedish subjects aged 50-65 years, a smaller cohort of 101 individuals were recruited for this study (named as ‘SCAPIS’) and were followed longitudinally for one year with repeated analyses at three months’ intervals. The study was approved by the Ethical Review Board of Gothenburg, Sweden. All participants have provided a written informed consent. The study was performed in accordance with the declaration of Helsinki. The subjects had been extensively phenotyped before entering the study.

#### Study design

This is a non-interventional and observational study with the aim of collecting longitudinal data in a community-based cohort. A blood sample was collected every third month (+/- 2 weeks) making a total of 4 visits per subject. A selection of questions from the initial SCAPIS questionnaire was repeated to note any changes in health and lifestyle factors between each visit such as infections, disease, medication, perceived health, and exercise level.

## METHOD DETAILS

### Sample preparation and storage

Blood drawn in heparinized tubes at 4 visits from each subject was spun at 2000 g for 10 min for collection of plasma. Following this, blood was diluted in PBS and PBMCs were isolated using density gradient centrifugation and were frozen at - 80^0^C in freezing medium containing 10% DMSO and 90% FBS until analysis by Mass cytometry.

### Antibody labelling

The panel of monoclonal antibodies used for this study are indicated in the Key Resources Table. Monoclonal antibodies were either purchased pre-conjugated from Fluidigm or in purified carrier/protein-free buffer formulation from other vendors. Purified antibodies were conjugated to lanthanide metals using the MAXPAR X8 polymer conjugation kit (Fluidigm Inc.), according to the manufacturer’s protocol. Antibody concentration before and after conjugation was measured by NanoDrop 2000 spectrometer (Thermo Fischer Scientific) at 280 nm. Following conjugation of antibodies, they were diluted 1:1 in Protein Stabilizer PBS (Candor Bioscience GmbH) prior to use in experiments.

### Thawing and resting of cells

Cryopreserved PBMCs were thawed in RPMI medium supplemented with 10% FBS, 1% penicillin streptomycin, and benzonase (Sigma-Aldrich). Cells were resuspended in RPMI medium supplemented with 10% FBS and 1% penicillin streptomycin and rested overnight at 37°C in 5% CO2 for cells to be revitalized. Samples were randomized in batches to avoid systematic technical variation due to PBMC sample processing (Brodin et al., 2019).

### Mass cytometry - staining samples using automation

Overnight rested cells were counted and checked for their viabilities. For live-dead discrimination, cells were stained with 2.5μM Cisplatin (Fluidigm Inc.) in RPMI without FBS for 5 min at room temperature, followed by quenching with RPMI containing FBS. Cells were then fixed with 1% formaldehyde (Polysciences Inc.), washed, and resuspended in CyFACS buffer (PBS with 0.1% BSA, 0.05% sodium azide and 2mM EDTA). For surface marker staining, cells were incubated for 30 min at 4°C with a 30ul cocktail of metal conjugated antibodies (Antibodies listed in Key Resources Table) targeting the surface antigens, washed with CyFACS and fixed with 4% paraformaldehyde, all of which performed using a custom built liquid handling robotic platform (Mikes et al., 2019).

### Mass cytometry - sample acquisition

Cells fixed in 4% paraformaldehyde were stained with Iridium-labelled DNA intercalator at a final concentration of 0.125 μM (MaxPar® Intercalator-Ir, Fluidigm Inc.) on the day of sample acquisition. Following incubation for 20 min at room temperature, cells were washed with CyFACS buffer twice, once with PBS, and twice with milliQ water. Cells were counted and diluted to 500,000 cells/ml in milliQ water containing 10% EQ Four Element Calibration Beads (Fluidigm Inc.) and filtered through a 35µm nylon mesh (Lakshmikanth and Brodin, 2019). Samples were acquired on one of two CyTOF2 mass cytometers (Fluidigm Inc.) using CyTOF software version 6.0.626 with noise reduction, a lower convolution threshold of 200, event length limits of 10-150 pushes, a sigma value of 3, and flow rate of 0.045 ml/min.

### Plasma protein quantification

Plasma protein data was generated from a total of 101 samples using a multiplex proximity extension assay (Olink AB, Uppsala) as described previously (Lundberg et al., 2011). For analysis, 20µL of plasma from each sample was thawed and sent to Olink AB for analysis. In this assay, plasma proteins are dually recognized by pairs of antibodies coupled to a cDNA-strand that ligates when brought into proximity by its target, extended by a polymerase and detected using a Biomark HD 96 3 96 dynamic PCR array (Fluidigm Inc.). Eleven Olink panels have been used as indicated in Key Resources Table, capturing a total of 978 proteins in each plasma sample.

## QUANTIFICATION AND STATISTICAL ANALYSIS

### Mass Cytometry Preprocessing and Gating

All FCS-files unrandomized using the CyTOF software (version 6.0.626) were transferred without any additional preprocessing. Files were normalized using our own in-house implementation of normalization software described previously (Finck et al., 2013). Cells populations were manually gated using a Brodin lab developed software, Cytopy. Cell frequencies were batch corrected using the ComBat algorithm provided in the sva R package.

### Self-Organizing Maps for Clustering of Mass Cytometry Data

For each lineage, up to 10 000 cells were subsampled (n = 393 files from 99 individuals) for clustering. Each parameter was Z-score transformed to account for variations in absolute intensity between channels, and the maximum intensity was set to the 99th percentile to remove extreme outliers. Samples were clustered using the som() function from the kohonen R package on a hexagonal 20x20 grid with rlen = 100. Hierarchical subclustering was performed and a cutoff was set to generate a final set of 10 clusters per cell lineage. Cluster frequencies were batch corrected using the ComBat algorithm provided in the sva R package.

### Aitchison’s distances

Aitchison distance metric was used given the compositional nature of cluster abundances (Aitchison, 1982). Distances among mass cytometry samples were calculated on the compositional vector of immune cell clusters as determined by Self-Organizing Maps. Distances were calculated using the aDist() function in the robCompostitions R package.

### Describing variation along a principal curve

To describe the variation of individual’s cell composition along a trend, we first induce a 2-dimensional Euclidean space by applying non-metric multidimensional scaling (MDS) method on distance matrix computed by aDist() function as described in above step. We used isoMDS() function from MASS R package. Each point in this 2-dimensional MDS space represents individual’ cell composition, and is projected to minimize the stress, the square root of the ratio of the sum of squared differences between the input distances. The basic idea is to project the variation among individual’s cell composition as captured by Aitchison’s distances by using distance preserving technique like MDS to 2-dimensional Euclidean space.

In the second step, we fitted a principle curve (Hastie and Stuetzle, 1989), which is a smooth one-dimensional curve that passes through the “middle” of a data provides a nonlinear summary of the data. The reason to fit a nonlinear trend to data is to capture the nonlinearity in compositional data distances i.e., small distances may have cell frequencies with large differences and continuum of variation among cell composition as pointed out by (Kaczorowski et al., 2017).

We used the algorithm implemented in R package princurve. The implementation uses first principal component of Principal Components Analysis (PCA) as the initialization, then fit a smooth curve by using cubic splines and local averaging to determine self-consistency.

### Estimating immune cell – protein associations

We performed independent run of a Bayesian Variable Selection Using Variational Method for each cell population from 32 cells populations as target variable and 378 protein variables as explanatory variables. A log linear regression model was fitted to find associations between a single target variable (cell population) and a set of explanatory protein variables. Using a Bayesian Variable Selection (BVS) approach (Carbonetto and Stephens, 2012), strength of association between target variable I.e. cell population and set of explanatory protein variables is estimated as the posterior probability of inclusion (PPI).

A selection of the significant associations can then be made by choosing those associations that have PPIs greater than a specific value, for example greater than 0.5 for the median probability model. We used two empirical based method to choose the threshold for detecting significant association among protein and cell populations: 1) By looking at histogram of PPI, where we set 0.99 as threshold to filter out most significant associations i.e., association with 1% or less than 1% chances of their exclusion from the model to predict the cell population. 2) by computing Bayesian false discovery rate (BFDR) threshold based on an a priori value of false discovery rate alpha. In our study, we set value of alpha equal to 0.01. In other words, we want a threshold on probabilities i.e. PPIs in such way that the probability of a type I error, or a false positive, should be less than 1%. We used BFDR formula implemented as part of R package DMBVS (Wadsworth et al., 2017). The estimated value of this BFDR is equal to 0.7553617 in our case. After applying BFDR threshold, we have 226 associations having PPI greater than BFDR out of total possible 378 * 32 = 12096 associations.

## DATA AND SOFTWARE AVAILABILITY

The datasets generated in the study are available upon request through a S3WP data committee which will ensure compliance with GDPR regulations. All software and code is freely available upon request to the authors.

## REFERENCES

Aguirre-Gamboa, R., Joosten, I., Urbano, P., van der Molen, R.G., van Rijssen, E., van Cranenbroek, B., Oosting, M., Smeekens, S., Jaeger, M., Zorro, M., et al. (2016). Differential Effects of Environmental and Genetic Factors on T and B Cell Immune Traits. Cell Reports 17, 2474–2487.

Alpert, A., Pickman, Y., Leipold, M., Rosenberg-Hasson, Y., Ji, X., Gaujoux, R., Rabani, H., Starosvetsky, E., Kveler, K., Schaffert, S., et al. (2019). A clinically meaningful metric of immune age derived from high-dimensional longitudinal monitoring. Nat Med, 25, 3.

Bernstein, H.B., Plasterer, M.C., Schiff, S.E., Kitchen, C.M., Kitchen, S., and Zack, J.A. (2006). CD4 Expression on Activated NK Cells: Ligation of CD4 Induces Cytokine Expression and Cell Migration. J Immunol 177, 3669–3676.

Brodin, P., and Davis, M. (2016). Human immune system variation. Nature Reviews Immunology 17, 21–29.

Brodin, P., Jojic, V., Gao, T., Bhattacharya, S., Angel, C., Furman, D., Shen-Orr, S., Dekker, C.L., Swan, G.E., Butte, A.J., et al. (2015). Variation in the Human Immune System Is Largely Driven by Non-Heritable Influences. Cell 160, 37–47.

Carbonetto, P., and Stephens, M. (2012). Scalable Variational Inference for Bayesian Variable Selection in Regression, and Its Accuracy in Genetic Association Studies. Bayesian Anal 7, 73–108.

Carr, E.J., Dooley, J., Garcia-Perez, J.E., Lagou, V., Lee, J.C., Wouters, C., Meyts, I., Goris, A., Boeckxstaens, G., Linterman, M.A., et al. (2016). The cellular composition of the human immune system is shaped by age and cohabitation. Nat Immunol 17, 461–468.

Casanova, J.-L., and Abel, L. (2015). Disentangling Inborn and Acquired Immunity in Human Twins. Cell 160, 13–15.

Damond, N., Engler, S., Zanotelli, V., Schapiro, D., Wasserfall, C.H., Kusmartseva, I., Nick, H.S., Thorel, F., Herrera, P.L., Atkinson, M.A., et al. (2019). A Map of Human Type 1 Diabetes Progression by Imaging Mass Cytometry. Cell Metab 29, 755–768.e5.

Dopico, X., Evangelou, M., Ferreira, R.C., Guo, H., Pekalski, M.L., Smyth, D.J., Cooper, N., Burren, O.S., Fulford, A.J., Hennig, B.J., et al. (2015). Widespread seasonal gene expression reveals annual differences in human immunity and physiology. Nature Communications 6, 7000.

Flores, G.E., Caporaso, G.J., Henley, J.B., Rideout, J., Domogala, D., Chase, J., Leff, J.W., Vázquez-Baeza, Y., Gonzalez, A., Knight, R., et al. (2014). Temporal variability is a personalized feature of the human microbiome. Genome Biol 15, 531.

Franceschi, C., Garagnani, P., Parini, P., Giuliani, C., and Santoro, A. (2018). Inflammaging: a new immune–metabolic viewpoint for age-related diseases. Nat Rev Endocrinol 14, 1–15.

Furman, D., Jojic, V., Kidd, B., Shen-Orr, S., Price, J., Jarrell, J., Tse, T., Huang, H., Lund, P., Maecker, H.T., et al. (2013). Apoptosis and other immune biomarkers predict influenza vaccine responsiveness. Molecular Systems Biology 9, 659–659.

Hägg, D.A., Olson, F.J., Kjelldahl, J., Jernås, M., Thelle, D.S., Carlsson, L., Fagerberg, B., and Svensson, P.-A. (2009). Expression of chemokine (C–C motif) ligand 18 in human macrophages and atherosclerotic plaques. Atherosclerosis 204, e15–e20.

Hastie, T., and Stuetzle, W. (1989). Principal Curves. J Am Stat Assoc 84, 502.

Horowitz, A., Strauss-Albee, D.M., Leipold, M., Kubo, J., Nemat-Gorgani, N., Dogan, O.C., Dekker, C.L., Mackey, S., Maecker, H., Swan, G.E., et al. (2013). Genetic and Environmental Determinants of Human NK Cell Diversity Revealed by Mass Cytometry. Sci Transl Med 5, 208ra145–208ra145.

Kaczorowski, K.J., Shekhar, K., Nkulikiyimfura, D., Dekker, C.L., Maecker, H., Davis, M., Chakraborty, A.K., and Brodin, P. (2017). Continuous immunotypes describe human immune variation and predict diverse responses. Proceedings of the National Academy of Sciences 114, E6097–E6106.

Klingberg, E., Oleröd, G., Konar, J., Petzold, M., and Hammarsten, O. (2015). Seasonal variations in serum 25-hydroxy vitamin D levels in a Swedish cohort. Endocrine 49, 800–808.

Krieg, C., Nowicka, M., Guglietta, S., Schindler, S., Hartmann, F.J., Weber, L.M., Dummer, R., Robinson, M.D., Levesque, M.P., and Becher, B. (2018). High-dimensional single-cell analysis predicts response to anti-PD-1 immunotherapy. Nat Med 24, 144.

Matzaraki, V., Kumar, V., Wijmenga, C., and Zhernakova, A. (2017). The MHC locus and genetic susceptibility to autoimmune and infectious diseases. Genome Biol 18, 76.

Nakaya, H.I., Wrammert, J., Lee, E.K., Racioppi, L., Marie-Kunze, S., Haining, N.W., Means, A.R., Kasturi, S.P., Khan, N., Li, G.-M., et al. (2011). Systems biology of vaccination for seasonal influenza in humans. Nature Immunology 12, 786–795.

Ngo, S.T., Steyn, F.J., and McCombe, P.A. (2014). Gender differences in autoimmune disease. Frontiers in Neuroendocrinology 35, 347–369.

Olin, A., Henckel, E., Chen, Y., Lakshmikanth, T., Pou, C., Mikes, J., Gustafsson, A., Bernhardsson, A., Zhang, C., Bohlin, K., et al. (2018). Stereotypic Immune System Development in Newborn Children. Cell 174, 1277–1292.e14.

Patin, E., Hasan, M., Bergstedt, J., Rouilly, V., Libri, V., Urrutia, A., Alanio, C., Scepanovic, P., Hammer, C., Jönsson, F., et al. (2018). Natural variation in the parameters of innate immune cells is preferentially driven by genetic factors. Nature Immunology 19, 302–314.

Piasecka, B., Duffy, D., Urrutia, A., Quach, H., Patin, E., Posseme, C., Bergstedt, J., Charbit, B., Rouilly, V., MacPherson, C.R., et al. (2018). Distinctive roles of age, sex, and genetics in shaping transcriptional variation of human immune responses to microbial challenges. Proceedings of the National Academy of Sciences 115, E488–E497.

Querec, T.D., Akondy, R.S., Lee, E.K., Cao, W., Nakaya, H.I., Teuwen, D., Pirani, A., Gernert, K., Deng, J., Marzolf, B., et al. (2008). Systems biology approach predicts immunogenicity of the yellow fever vaccine in humans. Nature Immunology 10, ni.1688.

Rao, D.A., Gurish, M.F., Marshall, J.L., Slowikowski, K., Fonseka, C.Y., Liu, Y., Donlin, L.T., Henderson, L.A., Wei, K., Mizoguchi, F., et al. (2017). Pathologically expanded peripheral T helper cell subset drives B cells in rheumatoid arthritis. Nature 542, 110.

Schutyser, E., Struyf, S., Menten, P., Lenaerts, J.-P., Conings, R., Put, W., Wuyts, A., Proost, P., and Damme, J. (2000). Regulated Production and Molecular Diversity of Human Liver and Activation-Regulated Chemokine/Macrophage Inflammatory Protein-3α from Normal and Transformed Cells. J Immunol 165, 4470–4477.

Shen-Orr, S.S., Furman, D., Kidd, B.A., Hadad, F., Lovelace, P., Huang, Y.-W., Rosenberg-Hasson, Y., Mackey, S., Grisar, F., Pickman, Y., et al. (2016). Defective Signaling in the JAK-STAT Pathway Tracks with Chronic Inflammation and Cardiovascular Risk in Aging Humans. Cell Systems 3, 374–384.e4.

Skundric, D.S., Cruikshank, W.W., Montgomery, P.C., Lisak, R.P., and Tse, H.Y. (2015). Emerging role of IL-16 in cytokine-mediated regulation of multiple sclerosis. Cytokine 75, 234–248.

Sobolev, O., Binda, E., O’Farrell, S., Lorenc, A., Pradines, J., Huang, Y., Duffner, J., Schulz, R., Cason, J., Zambon, M., et al. (2016). Adjuvanted influenza-H1N1 vaccination reveals lymphoid signatures of age-dependent early responses and of clinical adverse events. Nature Immunology 17, 204–213.

Templ, M., Hron, K., and Filzmoser, P. (2011). Compositional Data Analysis: Theory and Applications. Wiley 341–355.

Tsang, J.S., Schwartzberg, P.L., Kotliarov, Y., Biancotto, A., Xie, Z., Germain, R.N., Wang, E., Olnes, M.J., Narayanan, M., Golding, H., et al. (2014). Global Analyses of Human Immune Variation Reveal Baseline Predictors of Postvaccination Responses. Cell 157, 499–513.

Versteylen, M., Manca, M., Joosen, I., Schmidt, D., Das, M., Hofstra, L., Crijns, H., Biessen, E., and Kietselaer, B. (2016). CC chemokine ligands in patients presenting with stable chest pain: association with atherosclerosis and future cardiovascular events. Neth Heart J 24, 722–729.

Davis, M., and Brodin, P. (2018). Rebooting Human Immunology. Annual Review of Immunology 36, 1–22.

Vulcano, M., Albanesi, C., Stoppacciaro, A., Bagnati, R., D’Amico, G., Struyf, S., Transidico, P., Bonecchi, R., Prete, A., Allavena, P., et al. (2001). Dendritic cells as a major source of macrophage-derived chemokine/CCL22 in vitro and in vivo. Eur J Immunol 31, 812–822.

